# Intranasal *Fritillaria taipaiensis* nanovesicles alleviate acute lung injury via the lung-brain axis

**DOI:** 10.64898/2026.09.07.749870

**Authors:** Huanan Rao, Jin Pan, Jinxin Guo, Xiaominting Song, Shukun Gong, Tao Zhou, Qinghua Wu, Yan Huang, Wei Nie, Cheng Peng, Zheyuan Li, Chaoxiang Ren, Jin Pei

## Abstract

Acute lung injury (ALI) is a severe inflammatory syndrome frequently complicated by brain injury, which worsens outcomes. Here, we explored whether regulation of the lung-brain axis represents a therapeutic strategy for the management of ALI. We show that intranasal administration of *Fritillaria taipaiensis*-derived exosome-like nanovesicles (Ft-ELNs) ameliorates LPS-induced ALI and ALI-associated hypothalamic inflammation in mice. Ft-ELNs preferentially accumulate in the lungs, inhibiting TRPV1 in both the lungs and the hypothalamus to establish the lung-brain axis. In parallel, Ft-ELNs primarily regulate systemic humoral immunity by inhibiting the VEGFR1/PI3K/AKT/NFκB signaling pathway. *In vitro* experiments further confirmed that Ft-ELNs are efficiently internalized by alveolar epithelial MLE-12 cells, exerting protective effects by suppressing VEGFR1 expression. Mechanistically, we identified Ft-miR1 of Ft-ELNs as the key functional effector that directly targets the *Flt1* 3’UTR to inhibit VEGFR1 expression. These data suggest that Ft-ELNs attenuate lung-brain axis inflammation by targeting VEGFR1 and may hold potential as novel therapeutic agents for the treatment of severe ALI.

## Introduction

Acute lung injury (ALI) and its severe form, acute respiratory distress syndrome (ARDS), are major global public health challenges (Matthay et al., 2019; Thompson et al., 2017). ALI/ARDS affects about 10.4% of ICU patients, with nearly 3 million cases worldwide each year (Bellani et al., 2016; Parcha et al., 2021). In-hospital mortality is high, ranging between 35% and 46%, and exceeds 56% in severe cases. Additionally, ALI/ARDS is frequently accompanied by secondary brain dysfunction, particularly in the context of sepsis, which is associated with increased mortality and long-term cognitive impairment (Fan et al., 2018; Hanasono et al., 2013; Mazeraud et al., 2020). The pathophysiology of ALI involves uncontrolled systemic inflammation and multi-organ dysfunction driven by a cytokine storm, leading to alveolar-capillary barrier disruption, increased vascular permeability, pulmonary edema, and refractory hypoxemia (Ware and Matthay, 2000; Matthay and Zemans, 2011). Supportive ventilation remains the mainstay of ALI/ARDS management, underscoring the need for effective pharmacological therapies (Dellinger et al., 2013; Hotz et al., 2019).

In the early phase of ALI, pulmonary inflammatory signals reach the central nervous system (CNS) through neural and humoral pathways (Izumi et al., 2024; Chen et al., 2024; Xiao et al., 2025). Pulmonary vagal afferent fibers, particularly TRPV1-expressing nociceptors, detect injury-induced ROS and inflammatory mediators (IL-1β, IL-6, TNF-α), transducing these stimuli into action potentials and transmitting them to brainstem centers, which then rapidly relay signals to higher brain regions (Almanzar et al., 2025). This leads to systemic symptoms such as fever and anorexia, and causes neuroendocrine dysregulation via the hypothalamic-pituitary-adrenal axis (Ilanges et al., 2022; Geloso et al., 2024; Mehdi et al., 2025). In addition, the humoral pathway serves as a critical mechanism for spreading inflammatory mediators throughout the systemic circulation (Mazeraud et al., 2020). Specifically, following LPS challenge, alveolar macrophages, endothelial cells, and neutrophils release pro-inflammatory cytokines, chemokines, and damage-associated molecular patterns, which damage the alveolar-capillary barrier and enter the systemic circulation, thereby triggering systemic inflammatory response syndrome as well as CNS inflammation by disrupting the blood-brain barrier (Li et al., 2021; Gao and Hernandes, 2021; Xiao et al., 2025). Neural signaling may interact with and potentiate subsequent humoral inflammatory responses, forming a self-sustaining lung-brain inflammatory cycle (Saraiva-Santos et al., 2024). However, current ALI interventions remain largely lung-centered, highlighting the need to target the lung-brain inflammatory axis (Millar et al., 2024).

Chuanbeimu (*F. taipaiensis*) is a traditional Chinese medicinal herb widely used for respiratory disorders, with functions described in traditional Chinese medicine (TCM) as clearing heat, moistening the lungs, resolving phlegm, and relieving cough (Chinese Pharmacopoeia Commission, 2025). In TCM clinical practice, it is prescribed for syndrome patterns such as phlegm-heat obstructing the lung and qi-yin deficiency, which share certain symptom features with ALI, including dyspnea and collapse (termed “chuantuo” in TCM) (Zhang and Wu, 2017; Gao, 2007). Although the lung-protective effects of Chuanbeimu have been primarily attributed to its steroidal alkaloids, its exosome-like nanovesicles remain an underexplored bioactive component in ALI research (Du et al., 2020; Liu et al., 2022; Jin et al., 2022; Wang et al., 2022b; Huang et al., 2023). PELNs are nanosized vesicles carrying microRNAs (miRNAs), proteins, lipids, and other bioactive cargos, and are increasingly recognized as natural nanotherapeutics with cross-kingdom regulatory activity, excellent biocompatibility, low immunogenicity, and efficient cellular internalization (Wang et al., 2014; Mu et al., 2014; Dad et al., 2021; Teng et al., 2018; Cai et al., 2018). PELNs from several medicinal plants, including *Artemisia*-derived nanovesicles (ADNVs), *Platycodon grandiflorum* exosome-like nanoparticles (PGLNs), *Inula japonica* Thunb-derived exosome-like nanovesicles (INVs), Rehmanniae Radix-derived ELNs, have already demonstrated protective effects in ALI models (Ye et al., 2024; Fu et al., 2025; Tang et al., 2026; Qiu et al., 2023). Yet these studies have largely been confined to oral or intravenous administration, which may not fully exploit the potential of PELNs for direct respiratory delivery. Crucially, whether intranasally delivered PELNs can provide such coordinated lung-brain protection in ALI remains unclear.

MiRNAs, important bioactive cargos of PELNs, are small non-coding RNAs (∼20–25 nt) that regulate gene expression post-transcriptionally (Bartel, 2004; He and Hannon, 2004; Filipowicz et al., 2008; Jonas and Izaurralde, 2015). Upon loading into the RNA-induced silencing complex, miRNAs guide the complex to target mRNAs primarily through partial base-pairing of their 5′seed sequence with complementary sites, typically located in the 3′untranslated region (3′UTR), resulting in translational repression or mRNA decay (Lewis et al., 2005; Dad et al., 2021; Teng et al., 2018). Empirical research has demonstrated that miRNAs encapsulated within PELNs sourced from tea and *Panax ginseng* exert therapeutic effects via post-transcriptional gene regulation, highlighting their potential to mitigate inflammatory responses. This mechanism is further supported by the use of synthetic miRNA mimics, which replicate endogenous miRNA functions for sequence-specific gene silencing (Luo et al., 2025; Ma et al., 2024; Tembo et al., 2025).

Although Chuanbeimu is traditionally used for respiratory disorders, whether its exosome-like nanovesicles (Ft-ELNs) protect against ALI through the lung-brain axis remains unclear. Therefore, in this study, we isolated and characterized Ft-ELNs from *F. taipaiensis* and demonstrated that intranasal administration of Ft-ELNs alleviates ALI and secondary hypothalamic inflammation via neural and humoral pathways, namely, suppression of pulmonary TRPV1-mediated vagal nociceptive signaling and inhibition of the pulmonary VEGFR1 cascade. In addition, we identified Ft-miR1 as a key functional cargo of Ft-ELNs that directly targets *Flt1* to suppress VEGFR1 expression. These findings reveal a lung-brain axis-mediated mechanism underlying the protective effects of Ft-ELNs and highlight their translational potential as a nanotherapeutic for ALI.

## Results

### Ft-ELNs displayed exosome-like nanovesicle characteristics and preferentially accumulated in the lungs

The isolation workflow for Ft-ELNs is schematically depicted in **Fig 1A**. Briefly, Ft-ELNs were isolated from 20 g of dried *F. taipaiensis* bulbs and resuspended in 2 mL of DPBS. TEM revealed that Ft-ELNs exhibited the typical cup-shaped morphology and double-membrane ultrastructure characteristic of exosome-like vesicles (**Fig 1B**). NTA analysis showed Ft-ELNs as a uniform population with a mean particle diameter of 145.8 nm and a concentration of approximately 2.1 × 10¹⁰ particles/mL (**Figs 1C and S1**). Ft-ELNs displayed a negative surface charge of −43.46 mV (**Fig 1D**), indicating good colloidal stability. Analysis of Ft-ELNs cargo revealed nucleic acids primarily in the 100–250 bp range (**Fig 1E**) and identified four major protein bands spanning 20–98 kDa (**Fig 1F**). Notably, Ft-ELNs maintained total protein content of 16.94 ± 0.15 mg/mL over 9 months of storage at −80 °C **(Fig 1G**), underscoring their long-term stability. Together, these results support the further evaluation of Ft-ELNs as bioactive therapeutic candidates. To assess the biodistribution of Ft-ELNs in ALI mice, DiR-labeled Ft-ELNs were administered intranasally followed by *in vivo* and *ex vivo* imaging, with the experimental timeline depicted in **Fig 1H**. Importantly, pulmonary accumulation was detected at 2 h post-administration and increased progressively, with pronounced retention in the lungs confirmed by *ex vivo* imaging (**Figs 1I and 1J**).

**Figure 1.**
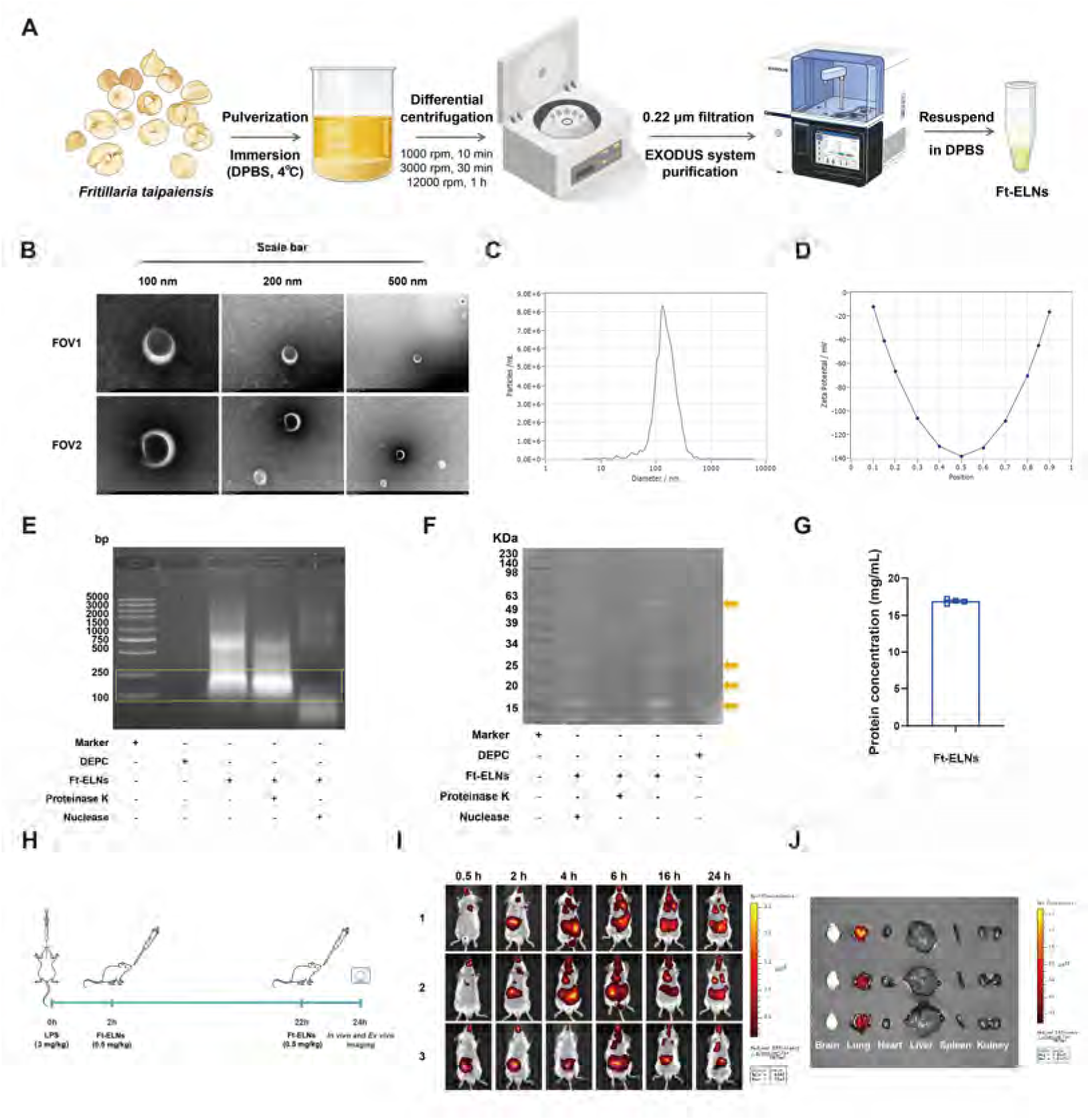
Characterization of FT-ELNs and their biodistribution in mice. **(A)** Schematic illustration of the isolation workflow for FT-ELNs from *F. taipaiensis* bulbs. **(B)** Representative TEM micrographs of Ft-ELNs from two fields of view (FOVs). Scale bars: 100, 200, and 500 nm. **(C)** Representative NTA results showing Ft-ELN size and concentration after a 100-fold dilution. **(D)** Zeta potential profile of Ft-ELNs. **(E)** Nucleic acids of Ft-ELNs were separated by 1.5% agarose gel electrophoresis. **(F)** Proteins of Ft-ELNs were visualized by 10% SDS-PAGE. **(G)** Storage stability of Ft-ELNs assessed by protein content. **(H)** Experimental timeline of intranasal administration and imaging for biodistribution analysis. **(I)** *In vivo* time-course imaging of DiR-labeled Ft-ELNs following intranasal administration. **(J)** *Ex vivo* imaging of excised organs.

### Ft-ELNs alleviated LPS-induced ALI through TRPV1 inhibition

Preliminary dose-ranging experiments identified 0.5 and 1mg/kg as the suitable intranasal doses (**Fig S2**), and this dose was used in all subsequent treatments. Experiments were conducted as outlined in **Fig 2A**. Micro-CT imaging demonstrated increased pulmonary attenuation and multifocal patchy opacities in LPS-treated mice, consistent with acute inflammatory lung injury and pulmonary edema. Ft-ELNs treatment alleviated these LPS-induced imaging abnormalities (**Fig 2B**). H&E staining corroborated these findings, confirming LPS-induced lung injury, as evidenced by alveolar septal thickening, inflammatory cell infiltration, interstitial edema, and disrupted lung architecture, which were markedly alleviated by Ft-ELNs treatment (**Figs 2C and 2D**). Notably, Ft-ELNs significantly inhibited LPS-induced TRPV1 upregulation in lung tissue (**Figs 2J and 2K**), suggesting suppression of TRPV1-mediated neurogenic inflammatory signaling. In line with this mechanism, immunohistochemical analysis revealed that Ft-ELNs markedly reduced the expression of the pro-inflammatory cytokines IL-1β, IL-6, TNF-α, and VEGF in lung tissue, indicating that Ft-ELNs effectively alleviated pulmonary inflammation and vascular hyperpermeability (**Figs 2E-2I**). In addition, Ft-ELNs reversed LPS-induced AQP5 downregulation, restored alveolar fluid clearance, and thereby attenuated pulmonary edema (**Figs 2L and 2M**).

**Figure 2.**
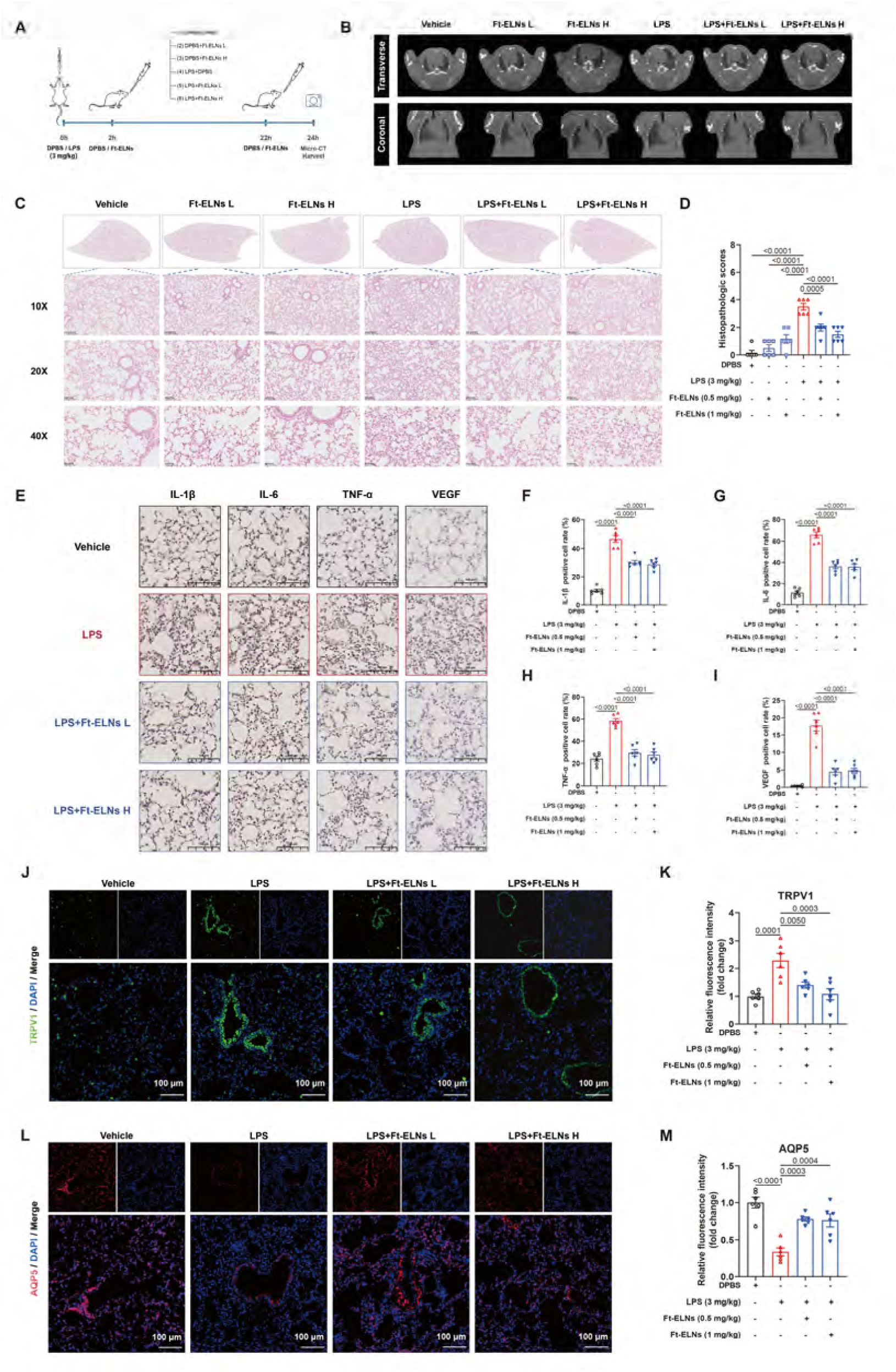
Therapeutic efficacy of intranasally administered Ft-ELNs in LPS-induced ALI mice. **(A)** Schematic of the *in vivo* experimental design and timeline. **(B)** Representative transverse and coronal *in vivo* micro-CT images of the lungs in mice. **(C)** Representative H&E-stained lung sections at ×10, ×20, and ×40 magnifications. **(D)** Histopathological lung injury scores among the six experimental groups. **(E)** Representative IHC staining of IL-1β, IL-6, TNF-α and VEGF in lung sections. Scale bars, 100 μm. **(F-I)** Semiquantitative analysis of IL-1β, IL-6, TNF-α and VEGF immunoreactivity. **(J)** Representative IF staining for TRPV1 in lung sections. Scale bars, 100 μm. **(K)** Semiquantitative analysis of TRPV1 immunofluorescence intensity. **(L)** Representative IF staining for AQP5 in lung sections. Scale bars, 100 μm. **(M)** Semiquantitative analysis of AQP5 immunofluorescence intensity. Data are presented as mean ± SEM (n = 6).

### Ft-ELNs alleviated ALI-associated hypothalamic and systemic inflammation

To assess whether ALI induces inflammatory injury in the brain, we examined coronal brain sections by H&E staining. LPS-challenged mice exhibited pronounced inflammatory cell infiltration in the hypothalamic paraventricular nucleus (PVN), which was markedly attenuated by treatment with Ft-ELNs (**Figs 3A and 3B**). Consistently, immunohistochemical analysis further revealed that Ft-ELNs effectively reversed the ALI-induced upregulation of IL-1β, IL-6, and TNF-α in the PVN (**Figs 3C and 3E-G**). These findings, together with the elevated TRPV1 expression observed in ALI mice, suggest that ALI triggered a TRPV1-driven neuroinflammatory response in the hypothalamus, which was effectively attenuated by Ft-ELNs treatment (**Figs 3J and 3K**). Moreover, ALI mice exhibited significantly increased VEGF expression and decreased Claudin-5 expression in the PVN, indicating BBB impairment in the hypothalamus. Ft-ELNs treatment largely reversed these changes (**Figs 3D, 3H and 3I**). In addition to ameliorating PVN inflammation and BBB-related alterations, Ft-ELNs significantly decreased the serum levels of IL-1β, IL-6, TNF-α, and endotoxin, and restored acetylcholine (ACh) levels in ALI mice, further indicating that Ft-ELNs mitigated systemic inflammatory responses (**Figs 3L–3P**).

**Figure 3.**
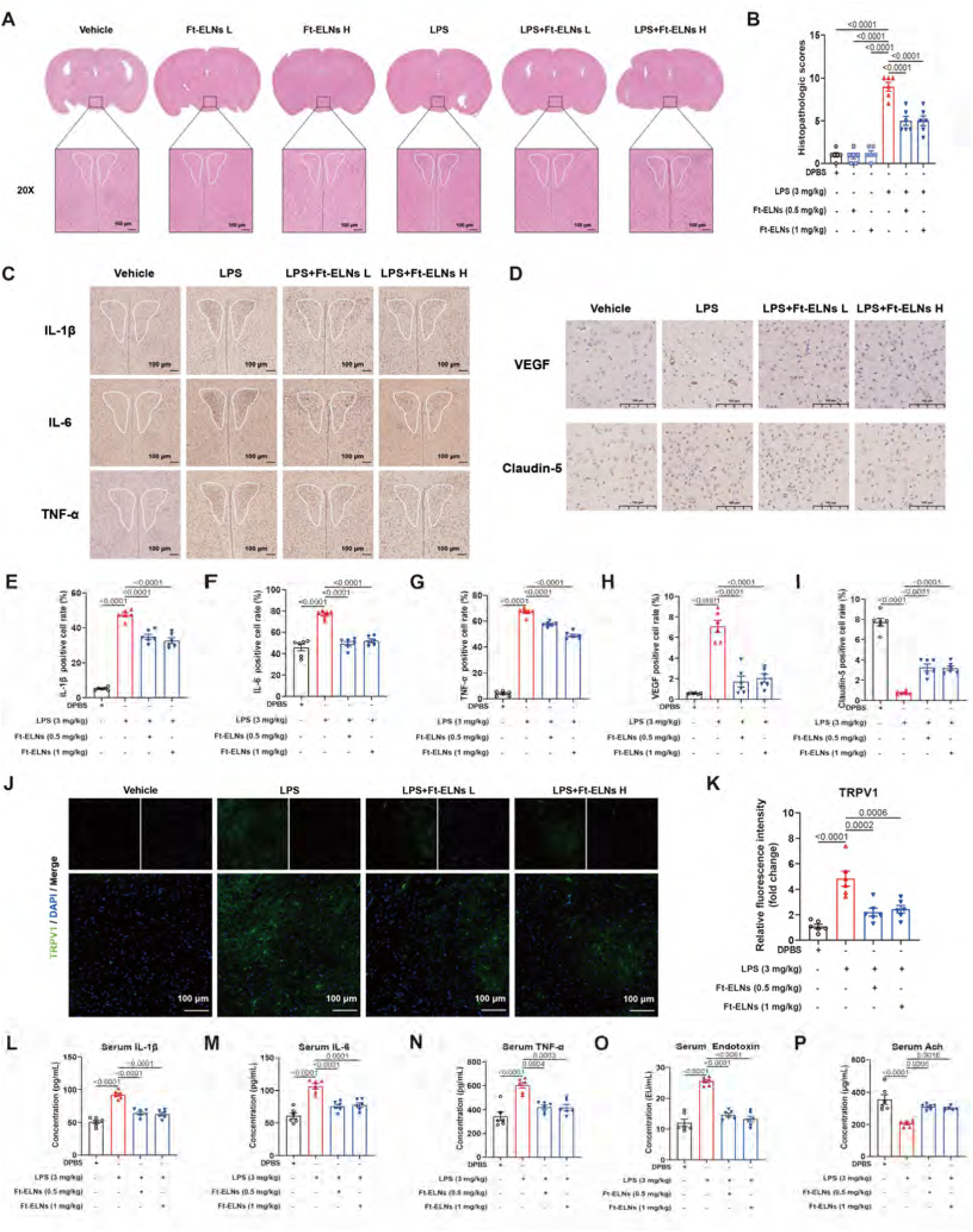
Therapeutic effects of Ft-ELNs against ALI-associated hypothalamic and systemic inflammation in mice. **(A)** Representative H&E-stained brain sections at ×20 magnification. **(B)** Histopathological PVN injury scores among the four experimental groups. **(C, D)** Representative IHC staining of IL-1β, IL-6, TNF-α, VEGF and Claudin-5 in the PVN region of brain sections. Scale bars, 100 μm. **(E-I)** Semiquantitative analysis of IL-1β, IL-6, TNF-α, VEGF and Claudin-5 immunoreactivity. **(J)** Representative IF staining for TRPV1 in the PVN region of brain sections. Scale bars, 100 μm. **(K)** Semiquantitative analysis of TRPV1 immunofluorescence intensity. **(L-P)** Serum IL-1β, IL-6, TNF-α, endotoxin and ACh levels. Data are presented as mean ± SEM (n = 6).

### Ft-ELNs suppressed the lung-hypothalamus inflammatory axis primarily through inhibition of VEGFR1/PI3K-AKT/NF-κB signaling

To investigate the molecular mechanisms underlying the protective effects of Ft-ELNs against ALI, we performed RNA-seq on mouse lung and hypothalamic tissues. The workflow for the lung transcriptome is shown in **Fig 4A**. Volcano plot analysis revealed that Ft-ELNs treatment induced 912 upregulated and 402 downregulated genes relative to the LPS group (**Fig 4B**). Venn analysis identified 371 shared genes among the WT, LPS, and LPS+Ft-ELNs groups, with 1788, 324, and 1292 unique genes in each group, respectively, indicating distinct transcriptional profiles **(Fig 4C**). GO and Reactome analyses showed that Ft-ELNs affected pathways related to defense response, innate immune regulation, ECM proteoglycans, and collagen chain trimerization, consistent with a shift from inflammatory to reparative processes (**Figs 4D and 4F**). KEGG analysis further revealed significant enrichment of the PI3K-AKT signaling pathway (**Fig 4E**). Given the important role of VEGFR1 in immune regulation, we next examined whether Ft-ELNs act through VEGFR1-mediated PI3K-AKT signaling. We found that Ft-ELNs reduced LPS-induced VEGFR1 expression, thereby decreasing the phosphorylation of PI3K, AKT, and NF-κB **(Fig. 4G and H**), thus protecting mice from ALI.

**Figure 4.**
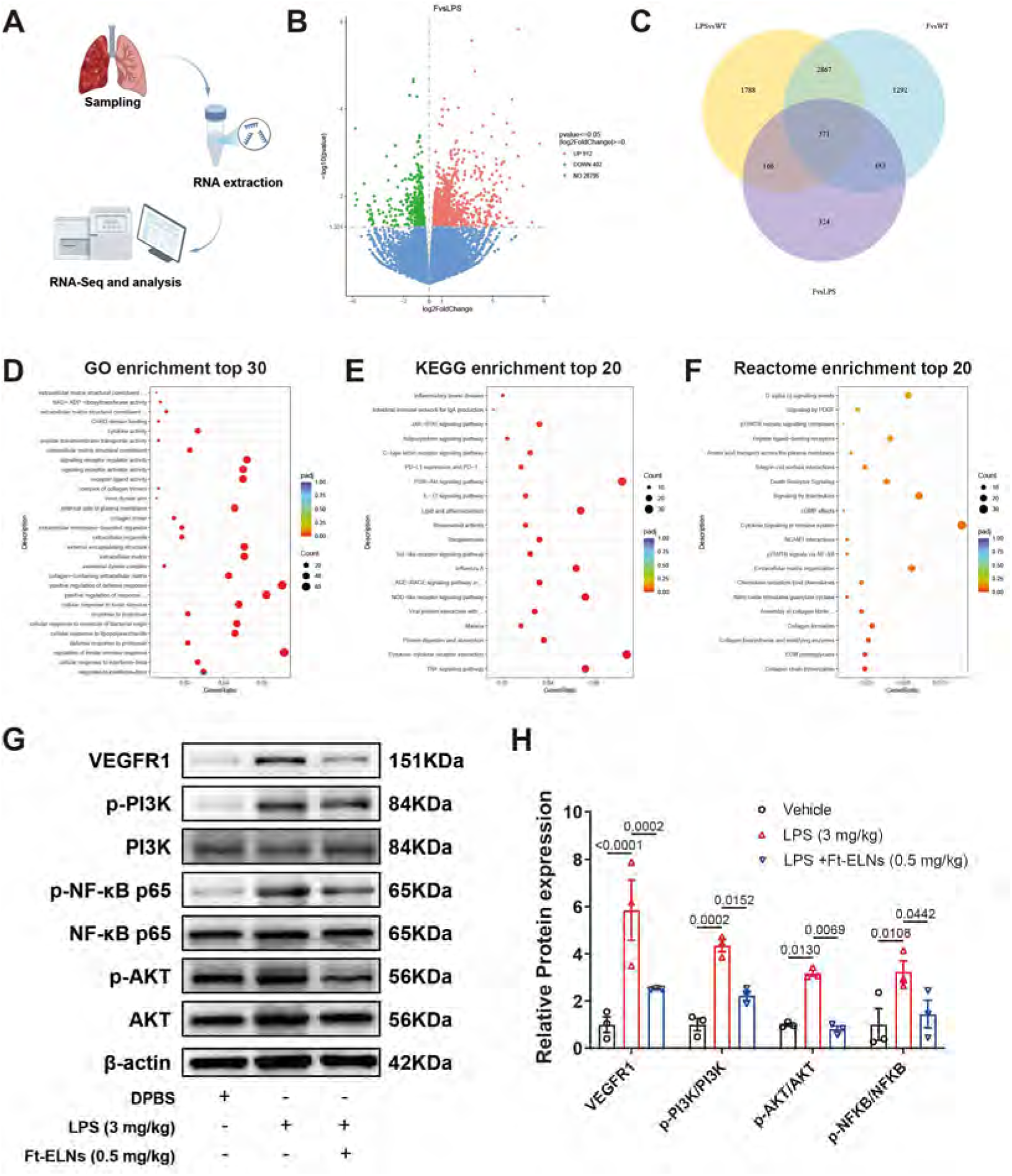
Transcriptomic and mechanistic analyses of Ft-ELNs-mediated protection in the lung. **(A)** Schematic of the workflow for lung tissue transcriptomic analysis. **(B)** Volcano plot of lung DEGs (LPS vs LPS+Ft-ELNs). **(C)** Venn diagram of lung DEGs across pairwise comparisons. **(D-F)** GO, KEGG, and Reactome enrichment analyses of lung DEGs (LPS vs. LPS+Ft-ELNs). **(G)** Representative Western blots of VEGFR1, p-PI3K, PI3K, p-AKT, AKT, p-NF-κB p65 and NF-κB p65 in mouse lung tissue. **(H)** Densitometric analysis of the blots. Data are presented as mean ± SEM (n = 3).

### Ft-ELNs attenuated LPS-induced injury in MLE-12 cells

Alveolar type II (AT2) epithelial cell dysfunction is a central pathological feature of ALI, driven by inflammatory disruption of the alveolar-capillary barrier and consequent pulmonary edema (Bi et al., 2025). To evaluate whether Ft-ELNs directly protect AT2 cells, we employed the MLE-12 cell line as an *in vitro* model (workflow in **Fig 5A**). CCK-8 assays showed that Ft-ELNs (0-100 μg/mL) were non-cytotoxic to MLE-12 cells after 48 h of treatment, as cell viability remained above 80% in both untreated and LPS (1 μg/mL)-stimulated cells (**Figs 5B and 5C**). High-content live-cell imaging further revealed dose-dependent cellular uptake of Ft-ELNs, with significant internalization observed at 10 μg/mL that was further augmented under LPS stimulation (**Figs 5D and 5E**). Functionally, Ft-ELNs significantly attenuated LPS-induced secretion of IL-1β, IL-6, and TNF-α, together with *Il6* and *Tnf* mRNA expression (**Figs 5F–J**), and concurrently reversed the upregulation of sVEGFR1 and VEGF (**Figs 5K and 5L**). These *in vitro* findings were consistent with *in vivo* observations, supporting the notion that Ft-ELNs directly act on pulmonary epithelial cells to exert anti-inflammatory effects and suppress vascular permeability factors.

**Figure 5.**
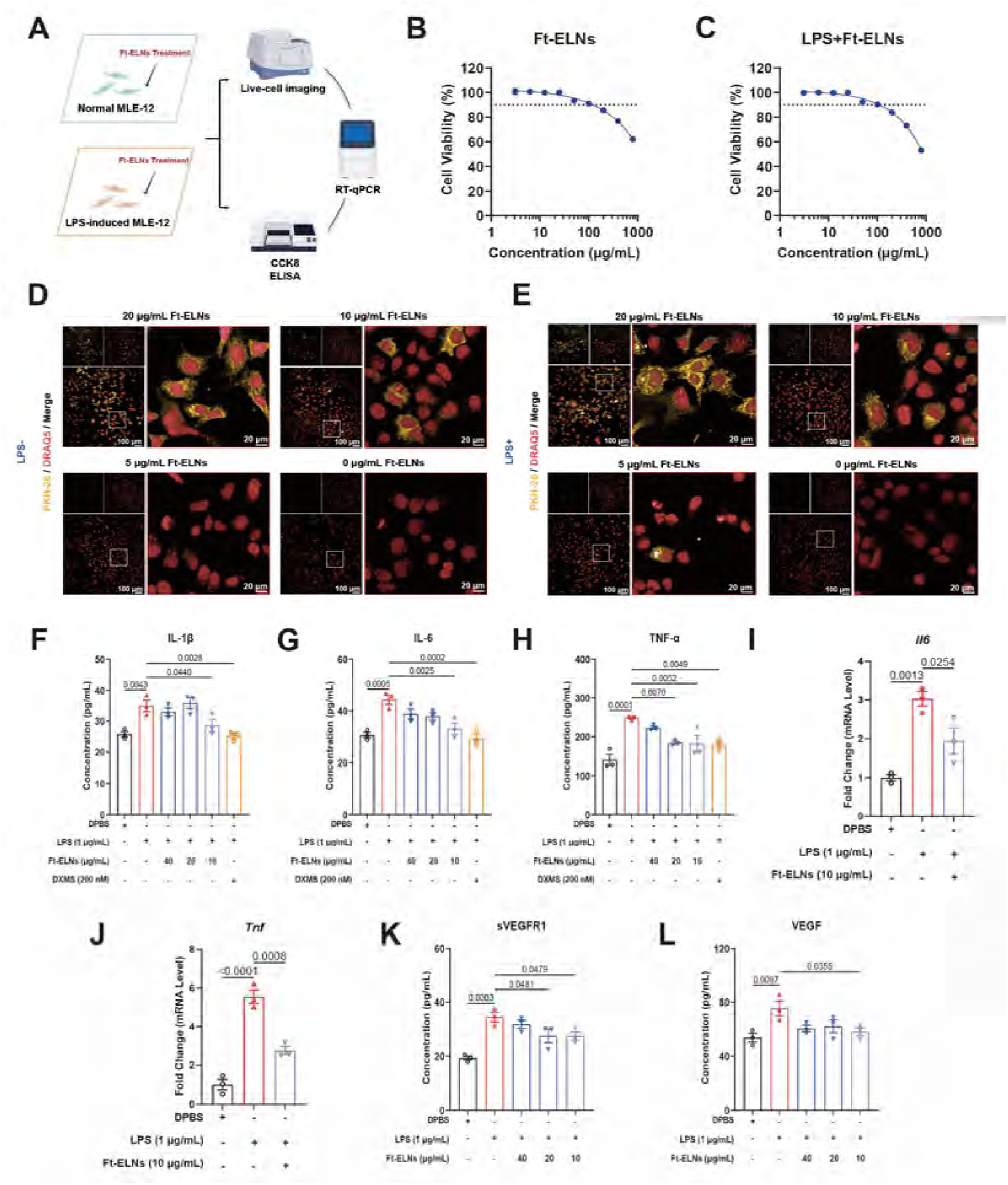
Internalization of Ft-ELNs and protection against LPS-induced injury in MLE-12 cells. **(A)** Schematic illustration of the experimental workflow. **(B, C)** Cell viability of MLE-12 cells treated with Ft-ELNs in the absence of LPS or presence of LPS (1 μg/mL). **(D, E)** Representative high-content images of live MLE-12 cells showing DRAQ5-stained nuclei (red) and PKH26-labeled Ft-ELNs (pseudocolored yellow) in the absence of LPS or presence of LPS (1 μg/mL). **(F-H)** Levels of IL-1β, IL-6 and TNF-α in cell culture supernatants. **(I-J)** RT-qPCR analysis of *Il6* and *Tnf* mRNA expression at 6 h. **(K-L)** Levels of sVEGFR1 and VEGF in cell culture supernatants. Data are presented as mean ± SEM (n = 3).

### De novo transcriptome assembly and functional annotation of *F. taipaiensis*

Total RNA from *F. taipaiensis* bulbs was sequenced, quality-filtered, and de novo assembled, and the resulting transcriptome dataset was annotated and analyzed (**Fig 6A**). Unigene lengths were predominantly distributed in the 501-1000 bp and 1001-2000 bp ranges, with the 501-1000 bp interval being the most abundant (**Fig 6B**). BUSCO assessment revealed that the whole transcriptome dataset contained 72.1% complete transcripts (55.4% single-copy, 16.7% duplicated), while the cluster and unigene datasets each showed 69.2% completeness (67.5% single-copy, 1.7% duplicated), with 11.2% fragmented and 19.6% missing transcripts (**Fig 6C**). A total of 4,098 sequences were annotated against the Nr, Nt, Pfam, KOG, and GO databases (**Fig 6D**). Among the annotated unigenes, 84.1% showed significant homology to Nr database sequences (E-value < 1e−30), 90.4% shared >60% similarity, and species matching revealed the highest homology to *Elaeis guineensis* (18.5%), followed by *Phoenix dactylifera* (17.4%) and *Cocos nucifera* (5.7%) (**Fig 6E**). GO annotation of 12907 unigenes revealed that’cellular process’ (biological process),’cellular anatomical structure’ and’protein-containing complex’ (cellular component), and’binding’ and’catalytic activity’ (molecular function) were the enriched terms (**Fig 6F**). KOG functional clustering indicated significant enrichment in transcriptional regulation, secretion, and transmembrane protein transport (**Fig 6G**). KEGG pathway analysis mapped 13232 unigenes to 31 major pathways across seven categories: Brite Hierarchies, Cellular Processes, Environmental Information Processing, Genetic Information Processing, Metabolism, Not Included in Pathway or Brite, and Organismal Systems (**Fig 6H**). Finally, MISA analysis of 25764 transcripts identified 2969 SSR loci, with trinucleotide repeats being the most frequent, followed by mononucleotide and dinucleotide repeats (**Fig 6I**).

**Figure 6.**
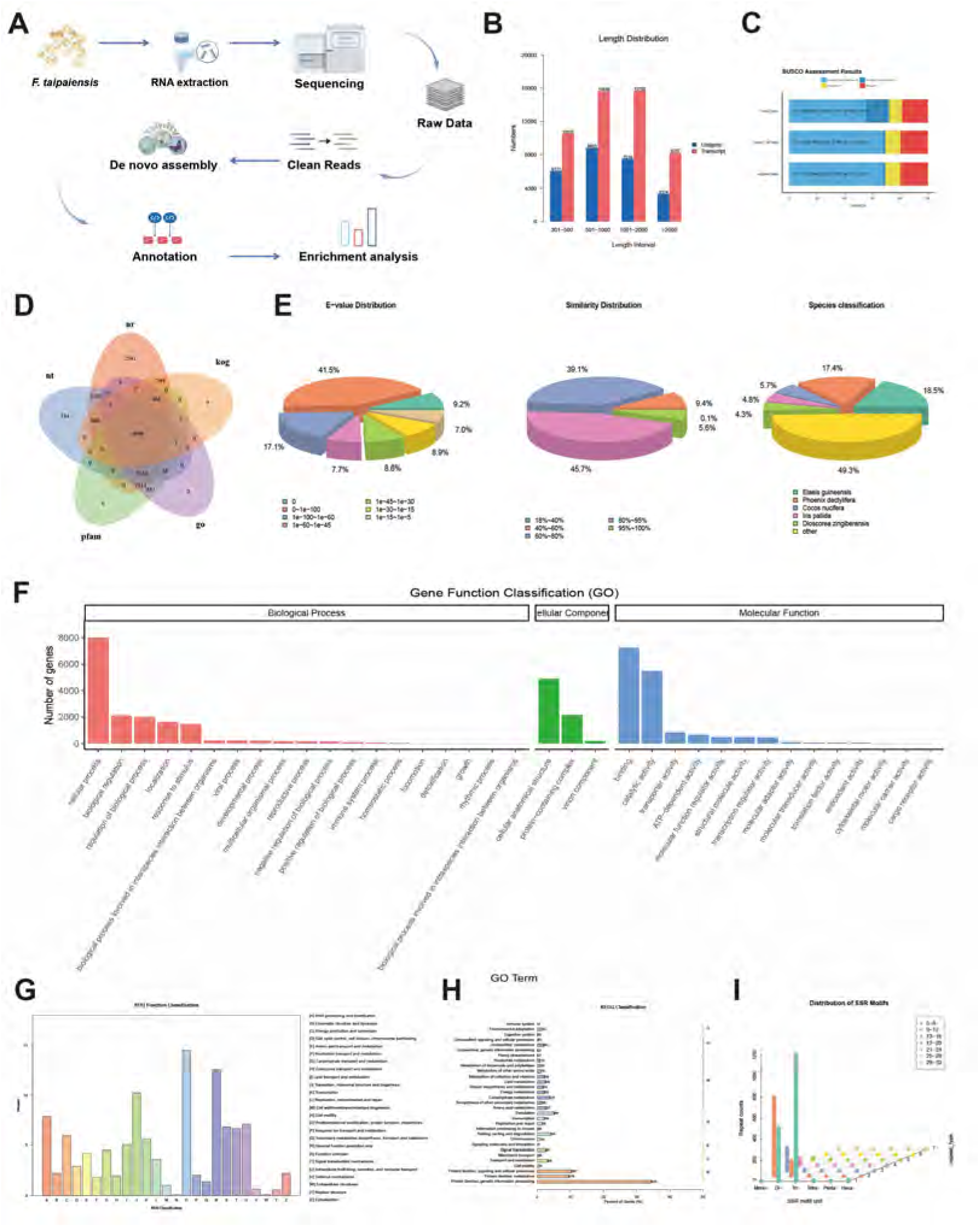
De novo transcriptome analysis of *F. taipaiensis*. **(A)** Schematic illustration of the overall workflow for de novo transcriptome analysis. **(B)** Length distribution of transcripts and unigenes. **(C)** BUSCO assessment of assembled transcripts. **(D)** Venn diagram of gene annotations from Nr, Nt, Pfam, KOG, and GO databases. **(E)** Statistics of sequence alignment against the NR database. **(F)** Statistics of GO annotation classification. **(G)** Statistics of KOG annotation classification. **(H)** Statistics of KEGG pathway classification. **(I)** Distribution of SSR density across transcripts.

### Identification and characterization of known and novel miRNAs in Ft-ELNs

Known and novel miRNAs in Ft-ELNs were analyzed as shown in **Fig 7A**. Clean reads were size-selected within the 18-30 nt range (**Fig 7B**), and the resulting sRNAs were mapped to the reference sequence and aligned against the *Arabidopsis thaliana* miRBase to identify known miRNAs. A total of 143 sRNA reads (25 unique) mapped to the reference genome, with 11 annotated as mature miRNAs. Known 21-nt miRNAs exhibited a first-position U preference. After filtering out novel miRNAs that aligned with known ncRNAs, three novel mature miRNAs were identified. Novel 21-nt miRNAs also preferred U at the first position, consistent with typical miRNA patterns (**Fig 7C**). In parallel, alignment against the *Oryza sativa* miRBase identified 216 sRNA reads (78 unique) mapped to the reference sequence, including 16 annotated mature miRNAs. Known 21-22 nt miRNAs similarly exhibited a first-nucleotide U preference. After excluding sequences matching known ncRNAs, three novel mature miRNAs were identified; however, novel 21 nt miRNAs showed a first-nucleotide C preference (**Fig 7D**). Altogether, 33 known and novel miRNAs were identified, and their secondary structures were predicted. Among the novel miRNAs, Ft-miR1 and Ft-miR5 were the most abundant (**Fig 7E**), and their predicted secondary structures are shown in **Fig 7F**.

**Figure 7.**
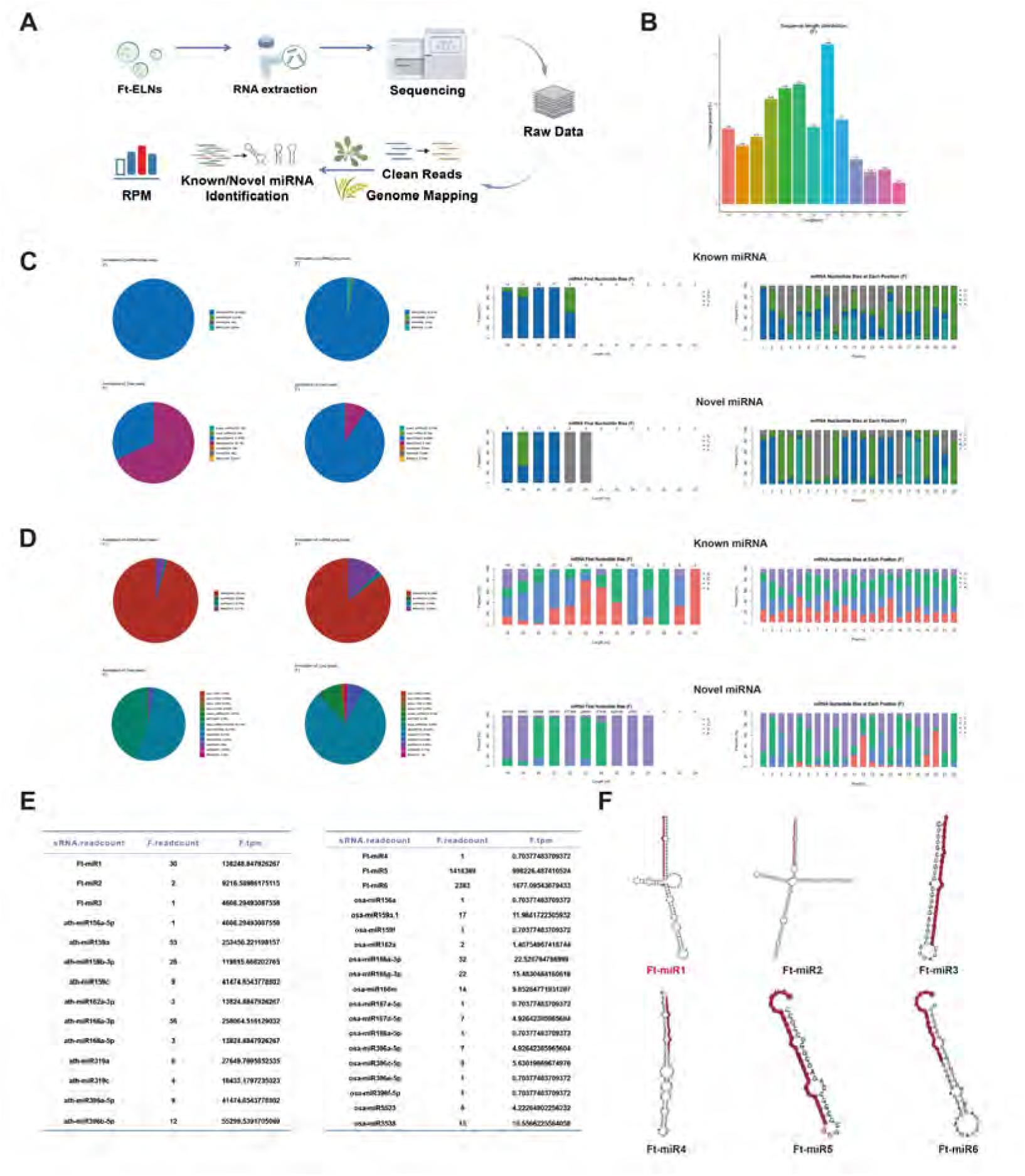
Small RNA sequencing analysis of Ft-ELNs. **(A)** Schematic illustration of the overall workflow for small RNA sequencing and analysis. **(B)** Length distribution of total sRNA fragment. **(C)** Annotation profiles of ncRNAs, miRNAs in total and unique small RNA reads, and nucleotide bias of known and novel miRNAs. The miRNAs were aligned to the miRBase with *Arabidopsis thaliana*. **(D)** Annotation profiles of ncRNAs, miRNAs in total and unique small RNA reads, and nucleotide bias of known and novel miRNAs. The miRNAs were aligned to the miRBase with *Oryza sativa*. **(E)** Expression profiling of mature miRNAs. **(F)** Secondary structures of novel miRNAs.

### Ft-miR1 attenuated LPS-induced injury by targeting *Flt1* 3ʹUTR in MLE-12 cells

Because Ft-miR1 better matched canonical miRNA characteristics, we transfected MLE-12 cells with the Ft-miR1 mimic to assess its function (**Fig 8A**). Ft-miR1 expression was significantly increased in the mimic-transfected group compared with the LPS control group, whereas no significant changes were observed in the Ft-miR1 inhibitor, mimic NC, or inhibitor NC groups, indicating successful transfection (**Fig 8B**). Western blot analysis showed that Ft-miR1 mimic transfection significantly reduced VEGFR1 expression, consistent with the effect observed after Ft-ELN treatment, whereas Ft-miR1 inhibitor transfection had no significant effect on its protein level (**Figs 8C and 8D**). Bioinformatics analysis identified two potential binding sites in close proximity within the *Flt1* 3’UTR (**Fig 8E**). A single fragment containing both binding sites was amplified and inserted into the pmirGLO dual-luciferase reporter vector to generate the wild-type (WT) reporter. The mutant (MUT) reporter, in which both binding sites were mutated, was then generated by site-directed mutagenesis (**Fig 8F**). Dual-luciferase reporter assays were then performed as detailed in **Fig 8G**. As shown in **Fig 8H**, co-transfection with Ft-miR1 significantly reduced the relative luciferase activity of the WT construct compared to the mimic NC, whereas the MUT construct showed no significant response. These results demonstrate that Ft-miR1 suppresses VEGFR1 expression at the post-transcriptional level by directly targeting the *Flt1* 3’UTR. Additionally, ELISA analysis revealed that Ft-miR1 significantly reduced LPS-induced IL-1β, IL-6, TNF-α, sVEGFR1, and VEGF levels in cell supernatants (**Figs 8I–8M**), suggesting that Ft-miR1 mediates the protective effects of Ft-ELNs against ALI *in vitro*.

**Figure 8.**
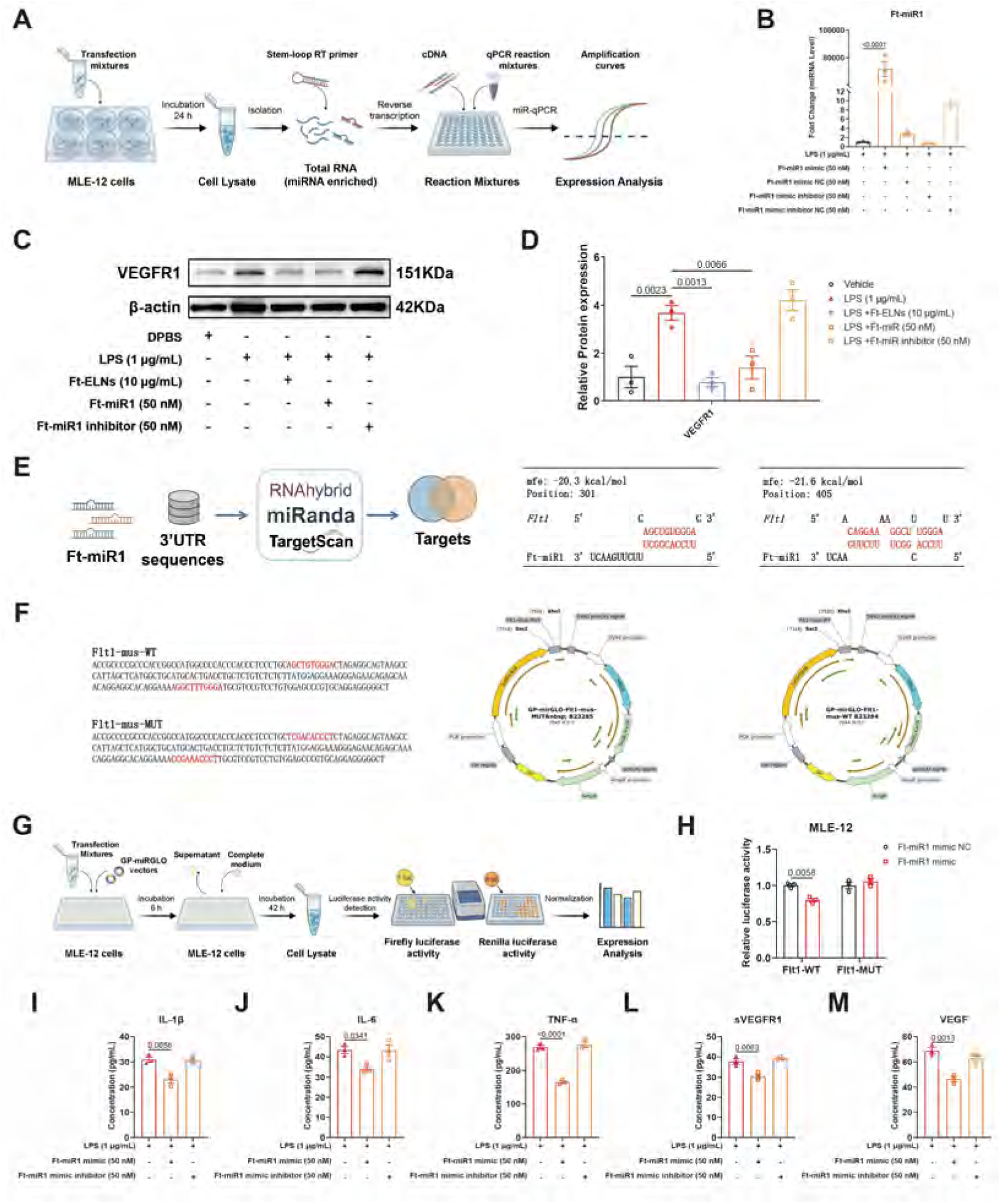
Validation of Ft-miR1 targeting the *Flt1* 3′UTR and suppressing VEGFR1 expression in MLE-12 cells. **(A)** Schematic illustration of the workflow for Ft-miR1 transfection efficiency validation. **(B)** Ft-miR1 expression measured by miR-qPCR and normalized to U6. **(C)** Representative Western blots of VEGFR1. **(D)** Densitometric analysis of the blots (n = 3). **(E)** Predicted high-affinity binding sites between Ft-miR1 and *Flt1* 3’UTR. **(F)** Schematic diagrams of the wild-type and mutant constructs. **(G)** Schematic illustration of the workflow for dual-luciferase reporter assay. **(H)** Dual-luciferase reporter assay of Ft-miR1 and 3’UTR of *Flt1*. Relative luciferase activity was calculated as the Firefly/Renilla ratio normalized to the NC group. **(I-M)** Levels of IL-1β, IL-6, TNF-α, sVEGFR1 and VEGF in cell culture supernatants. Data represent mean ± SEM (n = 3).

## Discussion

In this study, we isolated Ft-ELNs from *F. taipaiensis* and demonstrated their therapeutic efficacy against LPS-induced ALI. Ft-ELNs preferentially accumulated in the lungs after intranasal administration and effectively mitigated pulmonary histopathological injury, including inflammatory cell infiltration, alveolar septal thickening, and edema, while concurrently attenuating hypothalamic neuroinflammation and systemic inflammatory responses. Mechanistically, Ft-ELNs suppressed the lung-hypothalamus inflammatory axis primarily through inhibition of TRPV1 and VEGFR1/PI3K-AKT/NF-κB signaling. Furthermore, we identified Ft-miR1, a novel miRNA enriched in Ft-ELNs, which directly targets the *Flt1* 3’UTR to suppress VEGFR1 expression at the post-transcriptional level, thereby mimicking the anti-inflammatory effects of Ft-ELNs *in vitro*. These findings collectively position Ft-ELNs as promising plant-derived nanotherapeutics and suggest Ft-miR1 as a major mediator of their protective effects against ALI, highlighting a novel lung-brain strategy for integrated disease management and clinical translation.

Intranasal administration of PELNs overcomes several limitations associated with oral delivery, including gastric acid degradation, enzymatic degradation, and hepatic first-pass metabolism (Mu et al., 2023). As a non-invasive route, it enables rapid pulmonary deposition and accumulation, which may further modulate the lung-brain axis (Dantzer, 2018; Prescott et al., 2020). Although a fraction of the administered dose inevitably enters the gastrointestinal tract, direct pulmonary absorption is still expected to support rapid and targeted local bioavailability, with enhanced lung retention and reduced systemic exposure compared to intravenous administration (Djupesland, 2013; Fortuna et al., 2014; Qiao et al., 2021; Zhang et al., 2010). Importantly, intranasal delivery may achieve higher pulmonary exposure at lower doses than intravenous administration, thereby enhancing lung-targeting efficiency while reducing systemic distribution and off-target effects (Gou et al., 2025; Ahmed et al., 2025). These features suggest that intranasal delivery represents a promising therapeutic strategy for ALI, particularly in patients with hepatic impairment, by minimizing additional hepatic metabolic burden and potentially avoiding limitations associated with intravenous therapy (Damian et al., 2026).

Current standard care for ALI, primarily consisting of mechanical ventilation and systemic glucocorticoid therapy, remains limited by poor organ-targeting specificity, insufficient suppression of the systemic inflammatory cascade, and inadequate regulation of inter-organ crosstalk via conventional administration routes (Grotberg et al., 2023; Jenkins et al., 2024; Guan et al., 2023; Zhu et al., 2025). These limitations highlight the need for innovative adjunctive strategies, among which Ft-ELNs may represent a promising complement to existing interventions. Mechanical ventilation is indispensable for patients with ALI, yet it may paradoxically contribute to ventilator-induced lung injury (VILI) (Soumare et al., 2026). In this regard, Ft-ELNs may serve as a rapid therapeutic countermeasure against VILI by improving pulmonary barrier integrity and reducing microvascular permeability through the regulation of AQP5, Claudin-5, and VEGF. This protective effect may complement lung-protective ventilation strategies, thereby further alleviating pulmonary edema and severe hypoxemia. Systemic glucocorticoids are widely used to control inflammation in ALI; however, their clinical utility is frequently compromised by glucocorticoid resistance and dose-dependent adverse effects (Rouby and Lu, 2005). Ft-ELNs may help overcome this resistance by suppressing aberrant PI3K-AKT signaling, thereby potentially restoring glucocorticoid receptor sensitivity. Consequently, co-administration of Ft-ELNs with glucocorticoids may not only potentiate anti-inflammatory efficacy but also enable glucocorticoid dose reduction, thereby lowering the risk of complications such as infection and metabolic disturbances.

VEGFR1, a tyrosine kinase receptor for VEGF, is widely expressed in pulmonary microvascular endothelial cells, alveolar type II epithelial cells, and alveolar macrophages (Mura et al., 2006; Volpe et al., 2023; Domokos et al., 2023). Under pathological conditions, activation of the downstream PI3K/AKT signaling pathway via VEGFR1 exerts important regulatory effects in pulmonary fibrosis (Wang et al., 2022a; Amano et al., 2021). Our findings further suggest that during the early stage of ALI, VEGFR1 activation on inflammatory cells promotes pro-inflammatory PI3K/AKT signaling, thereby amplifying the inflammatory cascade and aggravating lung tissue injury. Given that persistent, uncontrolled inflammation is a major driver of the progression from ALI to pulmonary fibrosis (Liao et al., 2025), the anti-inflammatory effects of Ft-ELNs may have implications beyond the acute phase. By suppressing early inflammatory signaling—potentially through modulation of the VEGFR1/PI3K-AKT axis—Ft-ELNs may help limit the subsequent activation of pro-fibrotic pathways and delay fibrotic progression. Therefore, Ft-ELNs may represent a promising therapeutic strategy for both early-stage ALI and the prevention of post-ALI fibrotic remodeling.

We identified Ft-miR1 as a previously unrecognized component of Ft-ELNs and further demonstrated in MLE-12 cells that it suppresses VEGFR1 expression by directly targeting the *Flt1* 3′UTR, suggesting that this regulatory axis represents an important mechanism underlying the anti-ALI effects of Ft-ELNs. However, these findings are based on *in vitro* evidence, and the therapeutic efficacy of Ft-miR1 remains to be validated in ALI animal models. Moreover, given the diverse miRNA cargo harbored by Ft-ELNs, systematic screening and functional validation of additional miRNAs with anti-ALI activity are warranted. Such efforts may inform the design of multi-miRNA combination strategies, whether synergistic or sequential, to enhance therapeutic efficacy and expand the spectrum of clinical indications.

Although we demonstrated that Ft-ELNs effectively alleviate LPS-induced ALI and provided insight into the underlying molecular mechanisms, several limitations should be acknowledged. Most of the current evidence derives from experiments in the MLE-12 cell line and animal models, further preclinical toxicological evaluation and the development of scalable manufacturing protocols are required to support future clinical translation. Another limitation relates to the compositional complexity of Ft-ELNs. Beyond the functionally characterized Ft-miR1, the potential contributions and synergistic effects of other cargo components—including additional proteins and bioactive lipids—remain largely unexplored. Systematic integration of multi-omics analyses with functional rescue experiments will therefore be essential to define the individual roles of these components and to unravel their cooperative regulatory networks, thereby enabling a more comprehensive understanding of the pharmacodynamic basis of Ft-ELNs.

## Methods

### Isolation and characterization of Ft-ELNs

Ft-ELNs were isolated from the bulbs of *F. taipaiensis* (Wuxi County, Chongqing, China; botanically authenticated by Professor Jin Pei) using multi-step differential centrifugation. Briefly, dried bulbs were pulverized into fine powder and immersed in pre-cold DPBS at a 1:5 (w/v) ratio overnight at 4 °C. The mixture was centrifuged at 1000 rpm for 10 min, 3000 rpm for 30 min and finally 12,000 rpm for 1 h to remove cellular debris. The supernatant was filtered through a 0.22 μm filter (Millipore). Ft-ELNs were then purified from this supernatant using the EXODUS system (Huixin Lifetech, Shenzhen, China), resuspended gently with sterile DPBS. Samples were stored in aliquots at −80 °C until use. The protein content of Ft-ELNs were quantified using a BCA protein assay kit (A65453, Thermo Fisher Scientific, USA) immediately after extraction and quarterly thereafter (at months 3, 6, and 9) during 9 months.

The morphologies of Ft-ELNs were observed using a transmission electron microscope (TEM, Hitachi HT7700, Tokyo, Japan). The particle size, concentration and zeta potential of Ft-ELNs were measured by Nanoparticle tracking analysis (NTA) using ZetaView (PMX120, Particle Metrix, Germany). Ft-ELNs were lysed using an EV-specific lysis buffer (UR33101, Umibio, Shanghai, China). The supernatants of lysates were collected and treated with 100 μg/mL of Proteinase K (HY-108717, MCE, China) for 1 h at 55 °C or 50 U/mL of Nuclease (HY-131160, MCE, China) for 1 h at 37 °C. For nucleic acid analysis, agarose gel electrophoresis was performed using a 1.5% agarose gel at 120 V for 15 min in a 0.5× TBE buffer. The gel was stained with GelRed Nucleic Acid Stain (TSJ003, Tsingke, Beijing, China) and visualized using the Gel Doc XR+ (BioRad, Hercules, CA). For protein analysis, samples were boiled at 95 °C for 5 min with 1 × SDS loading buffer. A total of 22 μg per well of protein was separated on a SuperPAGE™ Precast Bis-Tris 10 % Gel (LK303, EpiZyme, China) at a constant voltage of 150 V for 40 min and imaged using transilluminated UV light with the ChemiDoc (Bio-Rad, Hercules, CA).

### Experimental animals

All animal experiments were approved by the Animal Ethics Committee of Chengdu University of Traditional Chinese Medicine (ethics no. 2026025), and all procedures adhered to the principles and guidelines of the National Institutes of Health. Male BALB/c mice (6–8 weeks old, 20−25 g) were obtained from Chengdu Dashuo Biotechnology Co., Ltd. [license number: SCXK-(Jing)2024−0001] and housed in specific pathogen-free (SPF) facilities with ad libitum access to standard chow and water. The animals were kept under controlled temperature (24 ± 2 °C), humidity (45 ± 10%), and light/dark conditions (12/12 h).

### Ft-ELNs *in vivo* and *ex vivo* imaging

Ft-ELNs suspension (1 mL) was gently mixed with 9 mL of a 100 μM DiR Iodide (40757ES25, YEASEN, Shanghai, China) prepared in sterile DPBS and incubated at 37 °C for 30 min in the dark. Following incubation, the free dye was removed using the EXODUS system, and the purified DiR-labeled Ft-ELNs were resuspended in 300 μL DPBS. The DiR-labeled Ft-ELNs (1.7 mg/mL) were then administered intranasally to LPS (3 mg/kg)-induced mice at a dose of 0.5 mg/kg. Bioluminescence imaging was performed *in vivo* at different time points post-administration and *ex vivo* at 24 h with an IVIS Spectrum imaging system (PerkinElmer, USA).

### Animal model establishment and treatment

The mice were randomly divided into six groups (6 mice/group): the vehicle (sterile DPBS), Ft-ELNs (0.5 mg/kg), Ft-ELNs (1 mg/kg), LPS (3 mg/kg), LPS + Ft-ELNs (0.5 mg/kg) and LPS + Ft-ELNs (1 mg/kg). Animals were anesthetized with isoflurane (4%/O2 gas for induction and 2%/O2 gas for maintenance; RWD Life Science, Shenzhen, China), followed by intratracheal administration of LPS (3 mg/kg) or an equivalent volume of DPBS, utilizing a MicroSprayer Aerosolizer (TOWINT Tech, Shanghai, China). At 2 h after modeling, Ft-ELNs or DPBS were administered via intranasal instillation. A second dose was given at 22 h post-modeling (2 h before euthanasia at 24 h).

### Micro-CT imaging *in vivo*

Micro-CT imaging was performed on anesthetized mice (2–5% isoflurane/O2 gas) using a Quantum GX micro-CT (PerkinElmer, Waltham, MA, USA). The scanning parameters included X-ray energy settings of 70 kV and 80 µA, a 4-min acquisition time, a field of view of 36 mm, and a resolution of 72 μm per pixel.

### Serum cytokines analysis

The mice were deeply anesthetized with 5% isoflurane. Blood samples were collected through retro-orbital bleeding and subsequently centrifuged (8,000 × g, 20 min, 4°C) to obtain serum before being euthanized. The serum levels of IL-1β, IL-6, TNF-α, endotoxin, and acetylcholine were quantified using the corresponding ELISA kits (Ruixin Biotech, Quanzhou, China) according to the manufacturer’s instructions. The kits used are listed in Supporting Information **Table S1**.

### Histopathological analysis

The left lungs and brain were perfused with 4% paraformaldehyde for 24 h. Paraffin sections (5 μm thick) were deparaffinized in xylene, rehydrated through graded ethanol series, and stained sequentially with hematoxylin (5 min) and eosin (5 min). After dehydration through ascending ethanol and clearing in xylene, slides were mounted with neutral balsam and digitized using a NanoZoomer S60 slide scanner (Hamamatsu Photonics, Japan). Histopathological sections were evaluated by two board-certified pathologists blinded to experimental groups using a semi-quantitative scoring system.

### Immunohistochemical (IHC)and immunofluorescence (IF) staining

For immunohistochemical staining, paraffin sections underwent dewaxing, antigen retrieval, endogenous peroxidase quenching, and serum blocking. They were then incubated with a primary antibody overnight at 4 °C, followed by a species-matched HRP-conjugated secondary antibody. The signal was developed with DAB and counterstained with hematoxylin before dehydration and mounting. For IF staining, paraffin sections were dewaxed, subjected to antigen retrieval, blocked with bovine serum albumin, and incubated overnight at 4 °C with primary antibodies. After incubation with fluorescence-conjugated secondary antibodies (protected from light), nuclei were counterstained with DAPI, autofluorescence was quenched, and slides were sealed with an anti-fluorescence quencher. All slides were imaged using the NanoZoomer S60 digital slide scanner. The antibodies used are listed in Supporting Information **Table S3**.

### Lung tissue transcriptome analysis

Total RNA was extracted from the right lung of mice using TRNzol Universal Reagent (DP424, TIANGEN, China) according to the manufacturer’s guidelines. RNA integrity (RIN) and concentration were assessed using an Agilent 2100 Bioanalyzer (Agilent Technologies, Santa Clara, USA). RNA-seq libraries were prepared using the Fast RNA-seq Lib Prep Kit V2 (RK20306, ABclonal, China) and sequenced on an Illumina NovaSeq 6000 PE150 platform by Novogene Co., Ltd. (Beijing, China). Raw reads were filtered with fastp, and clean reads were aligned to the Ensembl reference genome using HISAT2 (v2.2.1). Gene-level read counts were obtained with featureCounts (v2.0.6), and FPKM values were calculated based on gene length and read counts. Differential expression analysis was performed using R package DESeq2, and GO, KEGG, and Reactome enrichment analyses were conducted with clusterProfiler (v4.8.1).

### *F. taipaiensis* de novo transcriptome analysis

Total RNA was extracted from *F. taipaiensis* using TRNzol Universal Reagent. RNA integrity and concentration were evaluated with an Agilent 2100 Bioanalyzer. Libraries were quantified with a Qubit 2.0 fluorometer, diluted to 1.5 ng/μL, and further assessed by qPCR after insert size verification. Sequencing was performed on an Illumina NovaSeq 6000 PE150 platform by Novogene Co., Ltd. Raw reads were quality-checked with FastQC, trimmed to remove adapters and low-quality bases, and assembled de novo using Trinity. The resulting transcripts were clustered into components based on sequence similarity, and the longest isoform of each component was retained as the unigene for downstream annotation and enrichment analysis.

### Ft-ELNs miRNAs library construction and sequencing

Total RNA was extracted from Ft-ELNs using TRNzol Universal Reagent. After quality and quantity assessment, libraries were prepared using the TruSeq Small RNA Sample Prep Kit (Illumina). Libraries were diluted to 1.5 ng/μL based on Qubit 2.0 measurements, and insert size was verified using an Agilent 2100 Bioanalyzer. Libraries with the expected insert size were quantified by qPCR (effective concentration >2 nM), pooled, and sequenced on an Illumina SE50 platform by Novogene Co., Ltd.

### Ft-ELNs miRNAs analysis and cross-kingdom target prediction

Following quality control, raw small RNA sequencing reads were filtered to remove adapter contaminants, low-quality reads, low-complexity sequences, structural RNAs (rRNA, tRNA, snRNA, and snoRNA), and repetitive sequences. Clean sRNA reads (18-30 nt) were mapped to the de novo assembled reference sequence of *F. taipaiensis* using Bowtie. The annotation of known miRNAs was conducted by aligning their sequences with mature miRNAs cataloged in miRBase. *Arabidopsis thaliana* and *Oryza sativa* were employed as reference species owing to the comprehensive miRNA annotations. Candidate novel miRNAs were predicted using miREvo and miRDeep2 based on hairpin structure, Dicer cleavage signatures, and minimum free energy, followed by secondary structure evaluation using RNAfold. MiRNA expression levels were normalized to reads per million (RPM): RPM = (number of reads mapping to miRNA / number of reads in clean data) × 10^6^. To explore the potential cross-kingdom regulatory relevance of these miRNAs, putative mouse target genes were predicted against mouse 3’UTR sequences retrieved from Ensembl using RNAhybrid (v2.1.2) and miRanda (v3.3a). Only target sites meeting stringent criteria, including complementarity in the seed region (positions 2-8) and hybridization free energy <=-20 kcal/mol, were retained as a conservative filtering strategy. For downstream analysis, only targets consistently identified by both algorithms and additionally supported by TargetScan (v8.0) were retained.

### Cell culture

The mouse alveolar type II epithelial cells (MLE-12) were obtained from Zhong Qiao Xin Zhou Biotechnology Co., Ltd. (Shanghai, China) and maintained in the complete culture medium (ZM0470) provided by the company. Cells were cultured at 37 °C in a humidified incubator with 5% CO2, trypsinized using 0.25% trypsin (25200056, Thermo Fisher Scientific), and subcultured every 2 to 3 days depending on cell confluence.

### Ft-ELNs cellular uptake and internalization

Purified Ft-ELNs were labeled with 5 μM PKH26 (UR52302, Uimibio, Shanghai, China) at room temperature for 10 min, and unbound dye was removed using the EXODUS system. MLE-12 cells were seeded in 96-well plates at 5 × 10³ cells/well for overnight incubation. The cells were first exposed to LPS (1 μg/mL) or vehicle for 1 h, after which PKH26-labeled Ft-ELNs (5, 10, or 20 μg/mL) or free PKH26 dye were directly added without a medium change, followed by incubation for 24 h. Prior to imaging, cells were washed with DPBS, and nuclei were stained with 1 μM DRAQ5 (62254, Thermo Fisher Scientific) for 15 min. Live-cell imaging was performed using an Opera Phenix™ Plus High Content Screening System (Perkin Elmer, USA).

### Cell viability assay

MLE-12 cells were seeded in 96-well plates at a density of 5 × 10³ cells/well for overnight incubation. The cells were pre-treated with LPS (1 μg/mL) or vehicle for 1 h, followed by treatment with varying concentrations of Ft-ELNs for 48 h. The Ft-ELNs were administered in a 9-point, 2-fold serial dilution starting from a maximum concentration of 800 μg/ml. The inhibitory effect on cell proliferation was evaluated using a CCK-8 assay kit (BS350A, BioSharp, China).

### Secreted cytokines analysis

MLE-12 cells were seeded in 48-well plates at a density of 1.5 × 10^4^ cells/well for overnight incubation. The experimental groups were: vehicle (DPBS), LPS (1 μg/mL), LPS + Ft-ELNs (10, 20, or 40 μg/mL), and LPS + dexamethasone (DXMS, 200 nM, positive control). Cells in the co-treatment groups were pre-exposed to LPS for 1 h prior to the addition of Ft-ELNs or DXMS. After 24 h of incubation, supernatants were collected for ELISA analysis of IL-1β, IL-6, TNF-α, VEGF and VEGFR1 (Ruixin Biotech) according to the manufacturer’s protocols.

### RT-qPCR

MLE-12 cells were seeded at 1.5 × 10^5^ cells/well in 6-well plates for overnight incubation. The experimental groups were: the vehicle (DPBS), LPS (1 μg/mL), LPS + Ft-ELNs (10 μg/mL). After 6 h, total RNA was extracted using the SteadyPure RNA Extraction Kit (AG21024, Accurate Biology, Changsha, China), reverse-transcribed using HiScript III All-in-one RT SuperMix (R333-01, Vazyme, Nanjing, China), and subjected to qPCR using SupRealQ Ultra Hunter SYBR qPCR Master Mix (Q713-02, Vazyme). Relative fold changes were calculated using the comparative Ct method (2^−ΔΔCt), with *Gapdh* as the internal control. The primers used are listed in Supporting Information **Table S2**.

### Ft-miR1 mimic transfection and miRNA-qPCR

The miRNA mimic, inhibitor, and negative controls for Ft-miR1 were purchased from GenePharma (Shanghai, China). The mimic sequences were as follows: sense 5’-UUCCACGGCUUUCUUGAACU-3’, antisense 5’-UUCAAGAAAGCCGUGGAAUU-3’. To confirm transfection efficiency, MLE-12 cells were seeded at 1.5 × 10^5^ cells/well in 6-well plates and incubated overnight. Cells were then transfected with 50 nM Ft-miR1 mimic, Ft-miR1 inhibitor, Ft-miR1 mimic negative control (mimic NC), and Ft-miR1 mimic inhibitor negative control (inhibitor NC), respectively, using Hieff Trans® LipoBooster 3000 Transfection Reagent (40801ES01, YEASEN). After 24 h, small RNA-enriched total RNA was extracted using the SteadyPure RNA Extraction Kit. Reverse transcription of miRNA was performed using the miRNA First-Strand cDNA Synthesis Kit (AG11743, Accurate Biology) with stem-loop primers, followed by qPCR using the SupRealQ Ultra Hunter SYBR qPCR Master Mix. All miRNA qPCR primers were obtained from Accurate Biology. The relative expression level of Ft-miR1 was calculated using 2^−ΔΔCt, with U6 small nuclear RNA (snRNA) as the internal control. The primers used are listed in Supporting Information **Table S4**.

### Dual-luciferase reporter assay

The wild-type (WT) 3’UTR of *Flt1* and its mutant version (Mut, with site-directed mutagenesis of the Ft-miR1 binding site) were cloned into the GP-miRGLO vectors by GenePharma. To validate the direct targeting of Ft-miR1 on *Flt1*, MLE-12 cells were seeded in 96-well plates at a density of 1 × 10⁴ cells/well and incubated over night. Cells were then co-transfected with 100 ng of either the WT or Mut reporter plasmid, along with 50 nM Ft-miR1 mimic or mimic NC. After 6 h of transfection, the culture medium was replaced with complete medium. At 48 h post-transfection, luciferase activity was measured using the Dual-Luciferase Reporter Assay System (11402ES60, YEASEN) according to the manufacturer’s instructions on a Spectramax iD5 Multi-Mode Microplate Reader (Molecular Devices, USA). Relative luciferase activity was calculated as the ratio of Firefly to Renilla luciferase activity.

### Western blotting

Lung tissues were harvested and homogenized using a lapping instrument (SWE-FP PLUS, Servicebio, China) in RIPA lysis buffer (PC101, EpiZyme) supplemented with protease and phosphatase inhibitors (P1045, Beyotime, China). For cell experiments, MLE-12 cells were seeded at 1.5 × 10⁴ cells/well in 6-well plates and incubated overnight. Cells were divided into five groups: the vehicle group, LPS (1 μg/mL), LPS + Ft-ELNs (10 μg/mL), LPS + Ft-miR1 mimic (50 nM), and LPS + Ft-miR1 inhibitor (50 nM). After 24 h of incubation, cells were lysed in RIPA buffer supplemented with protease and phosphatase inhibitors on ice for 30 min. The lysates were centrifuged at 12,000 × g for 10 min at 4 °C, and the supernatants were collected for protein quantification using a BCA protein assay kit. Subsequently, 1× SDS loading buffer was added, and the mixture was boiled at 95 °C for 10 min.

Samples (20-30 μg/well) were separated by SDS-PAGE using gels prepared with the Omni-Easy™ One-Step PAGE Gel Fast Preparation Kit (PC222, EpiZyme) and 5× SWE Rapid High-Resolution Electrophoresis Buffer (G2152, Servicebio). The separated samples were subsequently transferred onto 0.45 μm PVDF membranes (G6047, Servicebio) at 300 mA for 60 min at 4 °C. The membranes were then immersed in TBST solution supplemented with 5% skim milk and blocked for 1 h at room temperature, followed by washing with TBST three times for 10 min each. The membranes were incubated with primary antibodies against VEGFR1, P-PI3K/PI3K, P-AKT/AKT, P-NFκB/NFκB, and β-actin overnight at 4 °C, followed by 1 h of incubation with secondary antibodiesat room temperature. Lastly, membranes were washed three times with TBST for 10 min each, and ECL luminescent solution (Abbkine, Wuhan, China) was added. Protein bands were then imaged using a chemiluminescence imager (SCG-W3000 PLUS, Servicebio).The antibodies used are listed in Supplementary Information **Table S5**.

## Statistical analysis

This study presented all data as mean ± SEM. ImageJ software was utilized for image analysis, and GraphPad Prism software (version 9.5.1) was used for all statishatical analyses. Parametric data were subjected to one-way ANOVA followed by the Tukey-Kramer test, whereas nonparametric data were subjected to the Kruskal-Wallis test followed by the Dunn test. Sample sizes for specific assays were described in the figure legends, and a *p-value*<0.05 was considered statistically significant.

## Data availability

The raw RNA-seq and miRNA-seq data generated in this study have been deposited in the NCBI Sequence Read Archive (SRA) database under BioProject accession numbers PRJNA1524071 (http://www.ncbi.nlm.nih.gov/bioproject/1524071), PRJNA1524320 (http://www.ncbi.nlm.nih.gov/bioproject/1524320), and PRJNA1524323 (http://www.ncbi.nlm.nih.gov/bioproject/1524323).

## Author contributions

Huanan Rao: Investigation, Methodology, Data curation, Formal analysis, Writing-original draft. Jin Pan: Investigation, Methodology, Data curation, Formal analysis. Jinxin Guo: Data curation, Investigation, Validation, Graphical Abstract drawing. Xiaominting Song: Investigation, Validation. Shukun Gong: Investigation, Resources. Tao Zhou: Formal analysis, Visualization. Qinghua Wu: Formal analysis, Data curation. Yan Huang: Investigation, Validation. Wei Nie: Resources, Validation. Cheng Peng: Supervision, Methodology. Zheyuan Li: Resources, Validation. Chaoxiang Ren: Conceptualization, Funding acquisition, Supervision, Writing-review & editing. Jin Pei: Conceptualization, Funding acquisition, Project administration, Supervision, Writing-review & editing.

## Acknowledgements

This work was supported by the Department of Science and Technology of Yunnan, China (Grant No. 202502AS100024). Thanks to Chen Sun from Chengdu University of TCM for his help in cell imaging.

---

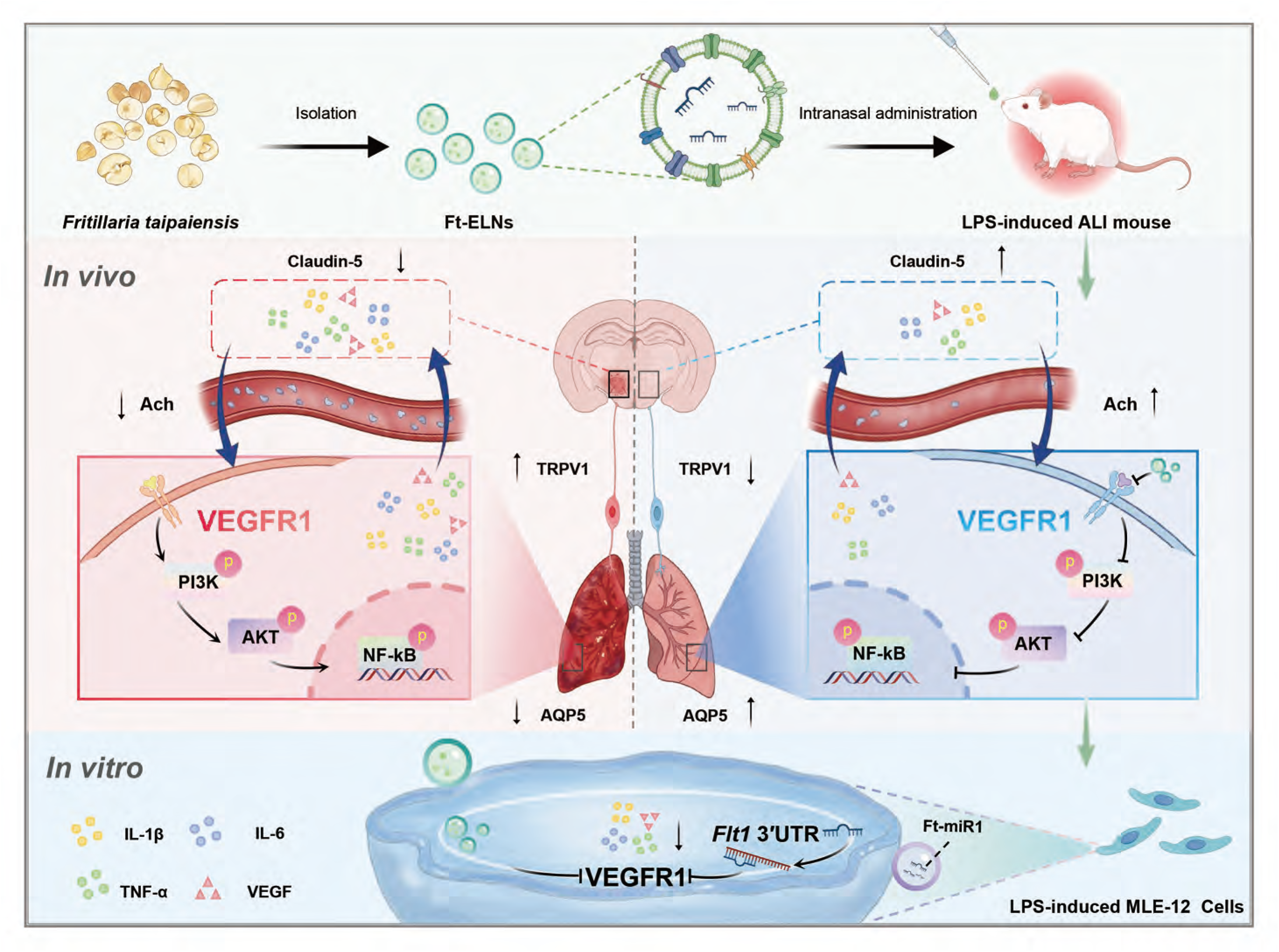

